# Factors associated with sharing email information and mental health survey participation in large population cohorts

**DOI:** 10.1101/471433

**Authors:** Mark J. Adams, W. David Hill, David M. Howard, Katrina A. S. Davis, Ian J. Deary, Matthew Hotopf, Andrew M. McIntosh

## Abstract

People who opt to participate in scientific studies tend to be healthier, wealthier, and more educated than the broader population. While selection bias does not always pose a problem for analysing the relationships between exposures and diseases or other outcomes, it can lead to biased effect size estimates. Biased estimates may weaken the utility of genetic findings because the goal is often to make inferences in a new sample (such as in polygenic risk score analysis). We used data from UK Biobank and Generation Scotland and conducted phenotypic and genome-wide association analyses on two phenotypes that reflected mental health data availability: (1) whether participants were contactable by email for follow-up) and (2) whether participants responded to a follow-up surveys of mental health. We identified nine genetic loci associated with email contact and 25 loci associated with mental health survey completion. Both phenotypes were positively genetically correlated with higher educational attainment and better health and negatively genetically correlated with psychological distress and schizophrenia. Recontact availability and follow-up participation can act as further genetic filters for data on mental health phenotypes.

## Introduction

Selection bias in epidemiological and cohort studies occurs when characteristics of individuals that influence their likelihood of becoming or remaining as study participants are also related to exposure to risk factors or to outcomes of interest (Hernán, Hernández-Díaz, & Robins, 2004). Selection bias can be introduced at many stages of a study, including at recruitment, at follow up, during record linkage, or in non-response to questionnaires or tasks and has the potential to lead to misestimates of phenotypic and genetic associations (Munafò, Tilling, Taylor, Evans, & Davey Smith, 2018). For example, a longitudinal study of psychiatric traits identified several characteristics related to loss-to-follow-up including age; education; ancestry; geographic location; and the presence, severity, and comorbidity of anxiety and depression (Lamers et al., 2012). There are several methods for handling selection bias if and when it needs to be taken into consideration. When all variables that influence selection and attrition are known, then bias can potentially be reduced or eliminated by conditioning on known variables or including them as predictors (Gelman & Hill, 2007). In longitudinal studies, techniques such as inverse probability weighting, where observations that are similar to those that were lost to follow-up contribute proportionally more to the analysis, can be used to correct for selection bias (Robins, Hernán, & Brumback, 2000). In study designs where the goal is to establish an association between an exposure and a disease outcome, selection bias is not an issue as long as there is sufficient variation in exposure (Fry et al., 2017).

Initial ascertainment and recontact have been demonstrated to have a genetic basis. For example, individuals who had a high genetic risk of schizophrenia (calculated from polygenic risk scores) were less likely to complete follow-up questionnaires or attend additional data collection sessions (Martin et al., 2016), and genetic liability for other traits have similar effects (Taylor et al., 2018). Participation in large cohort studies is already known to have a “healthy volunteer” effect (Fry et al., 2017) so we sought to characterise the phenotypic and genetic correlates of participation in follow-up studies that are focused on assessing mental health traits. To this end, we analysed recontact and participation in two studies: the Mental Health Questionnaire (MHQ) online follow-up in UK Biobank (Davis et al., 2018) and the Stratifying Resilience and Depression Longitudinally (STRADL) study in Generation Scotland (Navrady et al., 2018). We conducted phenotypic and genome-wide association analyses in UK Biobank to determine how participants who completed the MHQ differed from the rest of the sample. We also analysed factors related to whether UK Biobank participants were contactable by email, as email invitations were the primary method of recruitment into the MHQ follow-up. We used participation in the STRADL questionnaire follow-up in Generation Scotland as a replication data set for genetic findings.

## Methods

### Samples

UK Biobank (UKB) (Sudlow et al., 2015) is a population-based study of health in middle-aged and older individuals (N = 502,616). Eligible participants were aged 40 to 69 and recruited from 22 assessment centres in the United Kingdom. UK Biobank received ethical approval from the Research Ethics Committee (reference 11/NW/0382). The present study was conducted under UK Biobank application 4844.

Generation Scotland: Scottish Family Health Study (GS:SFHS) is a family-based cohort (N = 24,091) recruited through general practitioners in Scotland (Smith et al., 2012; Smith et al., 2006). Eligible participants were individuals aged 18 years or older who were able to recruit one or more family members into the study. GS:SFHS received ethical approval from the Tayside Research Ethics Committee (reference 05/S1401/89).

### Recontact and participation measures

During recruitment and baseline assessment (2006-2010), UKB participants were given the option of supplying an email address for receiving newsletters and invitations for online follow-up assessments. Of the 317,785 participants who supplied an email address, 294,738 provided a usable one while the remaining 23,047 either provided a syntactically incorrect or non-existent email address or asked that their email address be withdrawn. An email address was not provided by 184,831 UKB participants during baseline assessment. While this variable is called “email access” in the UK Biobank documentation (field 20005), we refer to this phenotype as “email contact”. Although additional UK Biobank participants have subsequently provided an email address for recontact (Davis et al., 2018), here we analyse the baseline availability of email contact so that it can be related to other baseline factors that were captured contemporaneously.

Starting in 2016, UKB participants who had provided email contact were sent an invitation to an online Mental Health Questionnaire (MHQ) entitled “thoughts and feelings” (Davis et al., 2018). Participants who had not started the questionnaire or had only partially completed it were sent reminder emails after two weeks and again after four months. Participants also received information about the MHQ in a postal newsletter with instructions on how to participate. From data supplied by UK Biobank on 12 June 2018, 157,396 participants had completed the MHQ. Responses to the MHQ were submitted between July 2016 and July 2017. Mean time between baseline assessment and MHQ follow-up was 7.5 years (range 5.9–11.2 years). We refer to this phenotype as “MHQ data”.

In 2015, GS:SFHS participants were sent a questionnaire package by post as part of the Stratifying Resilience and Depression Longitudinally (STRADL) project with the aim of studying psychological resilience (Navrady et al., 2018). Participants were eligible for follow up if they had consented to recontact and if they had a Community Health Index (CHI) number. Of the 21,525 eligible participants, 9,618 responded to the questionnaire, from which we coded a “STRADL data” phenotype.

### Phenotype analysis

Demographic and health differences between responders and non-responders to the STRADL survey have been analysed previously (Navrady et al., 2018) and found that, among other differences, participants who were women, non-smokers, or who had low levels of psychological distress were more likely to respond. We thus first conducted a similar analysis in UK Biobank. We ran logistic regressions for email contact and MHQ data using R 3.5.0 (R Development Core Team, 2018). We examined associations with age at initial assessment, sex, geographic region, educational qualification, smoking, alcohol consumption, number of diagnoses in linked electronic health records, and family history of dementia and depression.

We determined geographic region by grouping the assessment centres together into regions of England (South East, South West, East Midlands, West Midlands, North West, North East, and Yorkshire), Greater London, Scotland, and Wales. Education, smoking, drinking, and family history were assessed by means of a touchscreen interview during the initial assessment. We categorized educational qualifications as None, Professional, Higher (college or university), Secondary (A levels, O levels, GCSEs, CSEs), and Vocational (NVQ, HND, HNC). Smoking history had the responses ‘Prefer not to answer’, ‘Never’, ‘Previous’, and ‘Current’. For alcohol drinking, participants reported their average weekly and monthly consumption for different drink types from which we derived a measure of average alcohol consumption in units per week (Clarke et al., 2017) and standardized this variable for input into the model. For linked hospital records, we first removed diagnoses related to pregnancy (ICD-10 chapter O), congenital conditions (chapter Q), and health care provision (chapters U and Z). For the remaining diagnoses, we categorized them into mental health conditions and addictions (chapter F), injuries (chapter S, T, V, and Y), and all other diseases. We then counted the number of unique diagnostic codes each participant had for the three categories. Participants with linked hospital records who did not have any incidences of a diagnostic category were assigned a count of 0 while participants without linked records were set to missing.

### Genome-wide association, LD Score analysis, and replication analysis

UK Biobank contains genotype data imputed to ~92 million variants (Bycroft et al., 2017). We performed QC procedures on SNPs with filters for MAF > 0.001 and INFO > 0.1. We removed participants who had failed genotype platform QC, who did not cluster genetically as White British, or who overlapped with Psychiatric Genomics Consortium MDD and Generation Scotland participants; and we conducted additional filtering on related individuals (Howard et al., 2018). This resulted in 16,367,095 variants for 371,437 individuals for genetic analysis. We conducted genome-wide association analyses using BGENIE v1.3 (Bycroft et al., 2018) that coded the outcome variables as 0/1 in a linear regression. We covaried for age, sex, assessment centre, genotyping platform, and 20 UKB-provided principal components. We approximated odds ratios for the SNP effects using the transformation to the log-odds scale, log(OR) = *β* /(*k* (1 − *k*)), where *k* is the fraction of participants who were coded as 1 in the outcomes variable (email contact *k* = 0.6, MHQ data *k* = 0.33). We calculated SNP heritabilities on the liability scale using LD score regression (Bulik-Sullivan et al., 2015) and genetic correlations with 235 traits using LD Hub (Zheng et al., 2017). We used False Discovery Rate to correct for multiple testing when assessing the significance of the genetic correlations.

For Generation Scotland, 8,642,105 imputed variants were available for 19,994 participants (Hall et al., 2018). Variants with MAF < 0.005 and INFO < 0.8 were excluded. We performed association tests on the STRADL data phenotype using the mixed linear model with candidate marker excluded (MLMe) approach in GCTA v1.91.1 (Yang, Zaitlen, Goddard, Visscher, & Price, 2014). We constructed two GRMs using a leave-one-chromosome-out (LOCO) approach: one GRM that included all relationship coefficients and a second GRM that set relatedness to 0 when the relationship coefficients < 0.025 (Zaitlen et al., 2013). We fitted age and sex as covariates. To see if the results from the UKB phenotypes replicated, we looked up each independent significant SNP (or an LD proxy) in the GWAS of the STRADL data phenotype and assessed whether they were significant after Bonferroni correction. We also calculated the LD score genetic correlation of the STRADL data phenotype with the UKB email and MHQ data phenotypes.

### Loci discovery and functional annotation

Genomic risk loci were derived using clumping, carried out in FUnctional Mapping and Annotation of genetic associations (FUMA) (Watanabe, Taskesen, van Bochoven, & Posthuma, 2017). First, FUMA was used to identify independent significant SNPs using the *SNP2GENE* function. SNPs with a P-value of ≤ 5 × 10^−8^ and independent of other genome wide significant SNPs at r^2^ < 0.6 were identified from the summary GWAS statistics of the UKB email contact and MHQ data phenotypes. Second, using these independent significant SNPs, candidate SNPs were identified as all SNPs that had a MAF > 0.001 and were in LD of ≥ r^2^ 0.6 with at least one of the independent significant SNPs. These candidate SNPs included those from the UK10K/1000G and the haplotype reference consortia panel (UK Biobank release 1) and may not have been included in the UKB GWASs. Third, lead SNPs were identified using the independent significant SNPs. Lead SNPs were defined as SNPs that were independent from each other at r^2^ 0.1. Finally, genomic risk loci that were 250kb or closer were merged to form a single locus.

The lead SNPs identified above, and those in LD with the lead SNPs, were then mapped to genes using ANNOVAR and the Ensemble genes build 85. Intergenic SNPs were mapped to the two closest up and down stream genes which can result in them being assigned to multiple genes. eQTL mapping was performed using each independent significant SNP and those in LD with it. Those SNP-gene pairs that were not significant (FDR ≤ 0.05) were omitted from the analysis.

### Gene-mapping

Genetic variation in each of the independent genomic loci was mapped to genes using three complementary strategies. First, positional mapping was used to map SNPs to genes based on physical distance. SNPs within a 10kb window from the known protein genes found in the human reference assembly (hg19). Second, expression quantitative trait loci (eQTL) mapping was carried out by mapping SNPs to genes if allelic variation at the SNP was associated with expression levels of the gene. For eQTL mapping information on 45 tissue types from three data bases (GTEx, Blood eQTL browser, BIOS QTL browser) based on cis-QTLs where a SNPs are mapped to genes up to 1Mb away. A false discovery rate (FDR) of 0.05 was used as a cut off to define significant eQTL associations.

Finally, chromatin interaction mapping was carried out to map SNPs to genes when there is a three-dimensional DNA-DNA interaction between the independent genomic risk loci with a gene region. Chromatin interactions can involve long-range interactions between SNPs with genes as such no genomic distance boundary is applied. Hi-C data of 14 tissue types was used for chromatin interaction mapping. Chromatin interactions can also span multiple genes, and SNPs can be located in a region that interacts with other regions also containing multiple genes. In order to both reduce the number of genes mapped, and to increase the probability that those genes mapped are biologically linked to genetic variation at the independent genomic loci, only genes where one region involved with the interaction overlapped with a predicted enhancer region in any of the 111 tissue/cell types found in the Roadmap Epigenomics Project (Bernstein et al., 2010), and the other region was located in a gene promoter region (250bp upstream and 500bp downstream of the transcription start site and also predicted to be a promoter region by the Roadmap Epigenomics Project) were included here. An FDR of 1×10^-5^ was used to define a significant interaction.

### Gene-based GWAS

Gene-based analyses have been shown to increase the power to detect association due to the multiple testing burden being reduced, in addition to the effect of multiple SNPs being combined. Gene-based GWAS was conducted using MAGMA (de Leeuw, Mooij, Heskes, & Posthuma, 2015), also implemented in FUMA. Regardless of P-value, all SNPs located within protein coding genes were used to derive a P-value describing the association between genetic variation across the gene with either Email or questionnaire. The NCBI build 37 was used to determine the location and boundaries of 18,877 autosomal genes and linkage disequilibrium within and between genes was gauged using the UK Biobank 1 reference panel. A Bonferroni correction was applied to control for the number of genes tested.

### Gene-set analysis

A competitive gene-set analysis was conducted in MAGMA to identify the biological systems vulnerable to perturbation by common genetic variation. Competitive testing examines if genes within the gene set are more strongly associated with the trait of interest than genes from outside the gene set, and differs from self-contained testing by controlling for type 1 error rate as well as being able examine the biological relevance of the gene-set under investigation.

A total of 10,894 gene-sets (sourced from Gene Ontology, Reactome, and, MSigDB) were examined for enrichment. To control for the 10,894 gene sets examined, a Bonferroni correction was applied.

## Results

### Phenotypic associations of email contact and mental health follow-up (MHQ) data in UK Biobank

We conducted logistic regressions on email contact (valid Email address provided vs no valid Email address provided) and MHQ participation (those that had completed the MHQ vs those that had not completed the MHQ) in UK Biobank, examining the effects of age, sex, geographic region, educational attainment, drinking and smoking, and personal and family history of disease. We retained participants with complete data for analysis, which resulted in N = 294,381 for email contact (176,321 have email contact, 118,060 do not) and N = 294,381 for MHQ data (93,703 provided MHQ data, 200,678 did not). Odds ratios from the logistic regressions are listed in Table 1. Women in UK Biobank were less likely to have provided an email address for recontact but were more likely to be included in the MHQ. There was regional variation in email contact and MHQ data. Individuals who attended assessment centres in Greater London and the South West of England were the most likely to have provided an email address while individuals from assessment centres in the North East of England and Scotland were the least likely. Individuals with greater educational attainment, those who were not current smokers, those with a fewer number of hospital diagnoses, and those with a family history of dementia or severe depression were more likely to have email contact and to have MHQ data.

**Table 1.** Logistic regression on email contact (*N* = 294,381) and MHQ data (*N* = 294,381).

|  | Variable | Email contact |  | MHQ data |  |
| --- | --- | --- | --- | --- | --- |
|  |  | OR (SE) | P | OR (SE) | P |
| | Age | 0.98 (0.001) | $3.91 \times 10^{-281}$ | 1.00 (0.001) | 0.231 |
| Sex | Female | 1 | — | 1 | — |
| | Male | 1.12 (0.010) | $4.89 \times 10^{-37}$ | 0.90 (0.009) | $1.10 \times 10^{-32}$ |
| Region | East Midlands | 1 | — | 1 | — |
| | Greater London | 1.79 (0.042) | $7.59 \times 10^{-188}$ | 1.14 (0.020) | $3.78 \times 10^{-11}$ |
| | North East | 0.49 (0.010) | $1.47 \times 10^{-258}$ | 0.88 (0.018) | $2.00 \times 10^{-9}$ |
| | North West | 0.81 (0.014) | $8.28 \times 10^{-29}$ | 0.84 (0.015) | $4.05 \times 10^{-19}$ |
| | Scotland | 0.43 (0.010) | $< 2.23 \times 10^{-308}$ | 0.85 (0.017) | $6.61 \times 10^{-14}$ |
| | South East | 0.86 (0.018) | $9.78 \times 10^{-14}$ | 1.15 (0.025) | $3.02 \times 10^{-11}$ |
| | South West | 1.13 (0.026) | $3.36 \times 10^{-9}$ | 1.08 (0.023) | $3.39 \times 10^{-4}$ |
| | Wales | 0.59 (0.016) | $1.52 \times 10^{-106}$ | 0.84 (0.019) | $1.40 \times 10^{-12}$ |
| | West Midlands | 0.63 (0.013) | $4.66 \times 10^{-119}$ | 0.83 (0.017) | $2.69 \times 10^{-19}$ |
| | Yorkshire | 1.00 (0.021) | 0.86 | 0.94 (0.018) | $1.85 \times 10^{-4}$ |
| Qualifications | None | 1 | — | 1 | — |
| | Prefer not to answer | 1.01 (0.047) | 0.870 | 0.76 (0.050) | $1.78 \times 10^{-5}$ |
| | Higher | 4.28 (0.056) | $< 2.23 \times 10^{-308}$ | 4.42 (0.071) | $< 2.23 \times 10^{-308}$ |
| | Secondary | 2.66 (0.029) | $< 2.23 \times 10^{-308}$ | 2.61 (0.043) | $< 2.23 \times 10^{-308}$ |
| | Vocational | 2.06 (0.038) | $< 2.23 \times 10^{-308}$ | 2.07 (0.047) | $5.13 \times 10^{-241}$ |
| | Professional | 2.52 (0.045) | $< 2.23 \times 10^{-308}$ | 2.70 (0.063) | $< 2.23 \times 10^{-308}$ |
| Smoking | Prefer not to answer | 1 | — | 1 | — |
| | Never | 1.44 (0.107) | $2.32 \times 10^{-6}$ | 1.56 (0.144) | $1.18 \times 10^{-6}$ |
| | Previous | 1.63 (0.121) | $1.28 \times 10^{-10}$ | 1.64 (0.151) | $6.62 \times 10^{-8}$ |
|  | Current | 0.98 (0.074) | 0.639 | 1.07 (0.101) | 0.444 |
| Alcohol | Units/week (SD) | 1.04 (0.005) | $4.99 \times 10^{-18}$ | 1.02 (0.004) | $4.27 \times 10^{-6}$ |
| Diagnoses Yes vs No |  |  |  |  |  |
| | Mental disorder | 0.71 (0.020) | $3.58 \times 10^{-39}$ | 0.66 (0.022) | $1.13 \times 10^{-32}$ |
| | Injury | 0.92 (0.009) | $2.21 \times 10^{-23}$ | 0.91 (0.009) | $2.37 \times 10^{-24}$ |
| | Other disease | 0.97 (0.002) | $7.63 \times 10^{-99}$ | 0.93 (0.002) | $2.90 \times 10^{-286}$ |
| Family history Yes vs No |  |  |  |  |  |
| | Alzheimer's/dementia | 1.17 (0.013) | $3.68 \times 10^{-41}$ | 1.21 (0.016) | $5.81 \times 10^{-60}$ |
| | Severe depression | 1.05 (0.012) | $2.61 \times 10^{-5}$ | 1.12 (0.013) | $8.32 \times 10^{-23}$ |

### Genome-wide association analysis of email contact and MHQ data in UK Biobank

After filtering UK Biobank individuals to a White British, unrelated sample, the sample size was *N* = 371,417 for the GWAS of email contact and *N* = 371,428 for the GWAS of MHQ data. After clumping, there were nine loci (P ≤ 5 ×10^−8^) for email contact (Figure 1, Table 2, and Supplementary Table S1) and 25 for MHQ participation (Figure 2, Table 3, and Supplementary Table S11). The λ_GC_ was 1.29 for email contact and 1.37 for MHQ data.

**Table 2.** Top lead SNPs associated with email contact in UK Biobank (A1= effect allele, Freq. = frequency of A1 allele).

| Chr | SNP | Location (Bp) | A1/A2 | Freq. | OR (S.E.) | P-value |
| --- | --- | --- | --- | --- | --- | --- |
| 1 | rs632180 | 234,758,181 | T/C | 0.70 | 0.973 (0.005) | $2.0 \times 10^{-8}$ |
| 2 | rs7597665 | 34,420,702 | C/T | 0.29 | 1.031 (0.005) | $1.1 \times 10^{-9}$ |
| 2 | rs1455343 | 199,519,691 | T/G | 0.38 | 0.974 (0.005) | $2.2 \times 10^{-8}$ |
| 3 | rs73078357 | 48,695,834 | C/T | 0.12 | 1.038 (0.0066) | $4.5 \times 10^{-8}$ |
| 3 | rs111488606 | 49,864,924 | CA/C | 0.44 | 0.973 (0.005) | $2.3 \times 10^{-8}$ |
| 5 | rs6452788 | 87,712,913 | A/G | 0.24 | 1.032 (0.0054) | $2.9 \times 10^{-9}$ |
| 5 | rs4976602 | 167,843,998 | A/G | 0.11 | 0.96 (0.0069) | $2.7 \times 10^{-8}$ |
| 6 | rs1487441 | 98,553,894 | A/G | 0.49 | 1.031 (0.0047) | $9.5 \times 10^{-12}$ |
| 18 | rs1788784 | 21,159,630 | G/A | 0.66 | 1.031 (0.0042) | $1.3 \times 10^{-10}$ |

**Table 3.**
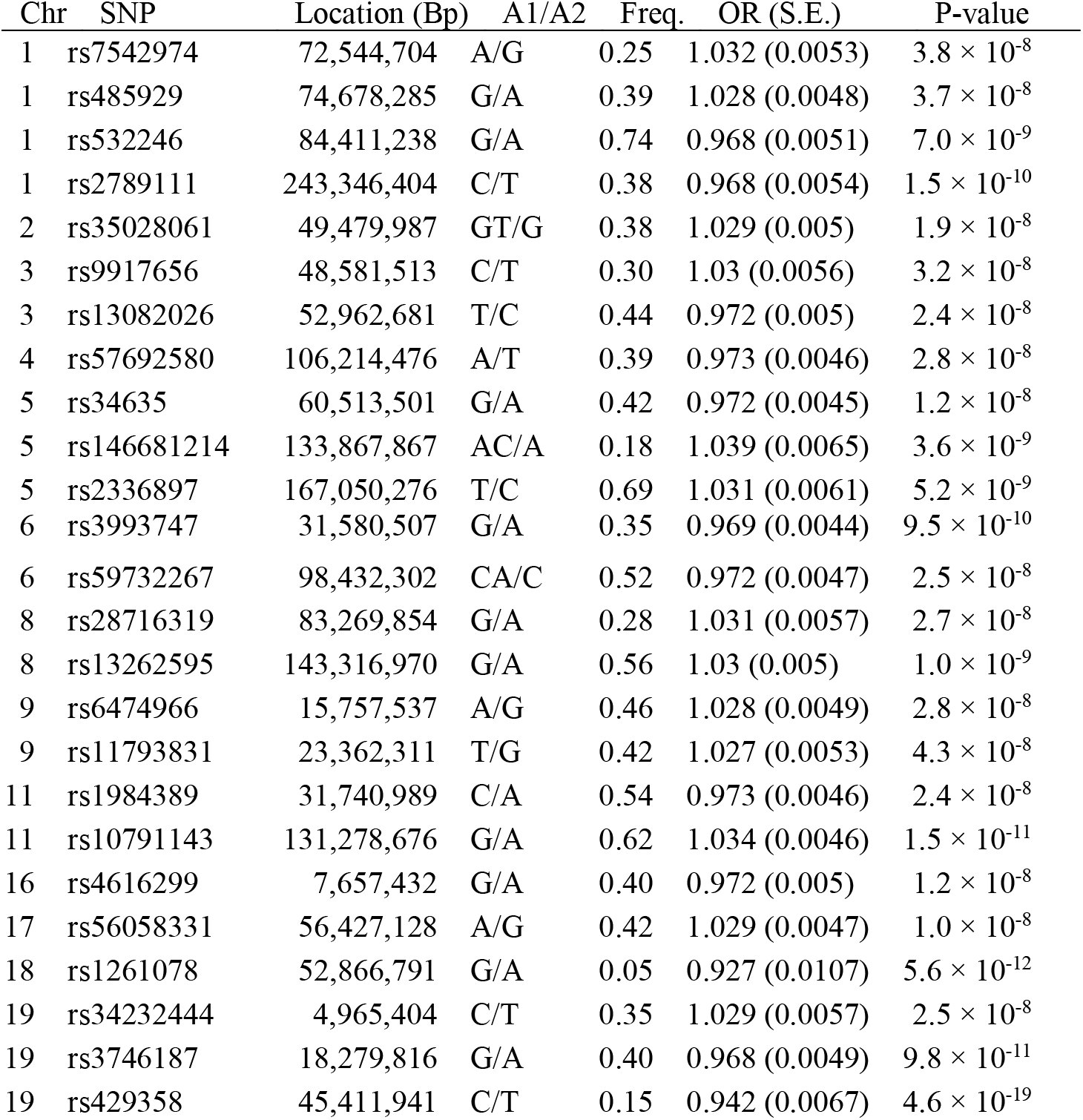
Top lead SNPs associated with MHQ data (A1= effect allele, Freq. = frequency of A1 allele.

| Chr | SNP | Location (Bp) | A1/A2 | Freq. | OR (S.E.) | P-value |
| --- | --- | --- | --- | --- | --- | --- |
| 1 | rs7542974 | 72,544,704 | A/G | 0.25 | 1.032 (0.0053) | $3.8 \times 10^{-8}$ |
| 1 | rs485929 | 74,678,285 | G/A | 0.39 | 1.028 (0.0048) | $3.7 \times 10^{-8}$ |
| 1 | rs532246 | 84,411,238 | G/A | 0.74 | 0.968 (0.0051) | $7.0 \times 10^{-9}$ |
| 1 | rs2789111 | 243,346,404 | C/T | 0.38 | 0.968 (0.0054) | $1.5 \times 10^{-10}$ |
| 2 | rs35028061 | 49,479,987 | GT/G | 0.38 | 1.029 (0.005) | $1.9 \times 10^{-8}$ |
| 3 | rs9917656 | 48,581,513 | C/T | 0.30 | 1.03 (0.0056) | $3.2 \times 10^{-8}$ |
| 3 | rs13082026 | 52,962,681 | T/C | 0.44 | 0.972 (0.005) | $2.4 \times 10^{-8}$ |
| 4 | rs57692580 | 106,214,476 | A/T | 0.39 | 0.973 (0.0046) | $2.8 \times 10^{-8}$ |
| 5 | rs34635 | 60,513,501 | G/A | 0.42 | 0.972 (0.0045) | $1.2 \times 10^{-8}$ |
| 5 | rs146681214 | 133,867,867 | AC/A | 0.18 | 1.039 (0.0065) | $3.6 \times 10^{-9}$ |
| 5 | rs2336897 | 167,050,276 | T/C | 0.69 | 1.031 (0.0061) | $5.2 \times 10^{-9}$ |
| 6 | rs3993747 | 31,580,507 | G/A | 0.35 | 0.969 (0.0044) | $9.5 \times 10^{-10}$ |
| 6 | rs59732267 | 98,432,302 | CA/C | 0.52 | 0.972 (0.0047) | $2.5 \times 10^{-8}$ |
| 8 | rs28716319 | 83,269,854 | G/A | 0.28 | 1.031 (0.0057) | $2.7 \times 10^{-8}$ |
| 8 | rs13262595 | 143,316,970 | G/A | 0.56 | 1.03 (0.005) | $1.0 \times 10^{-9}$ |
| 9 | rs6474966 | 15,757,537 | A/G | 0.46 | 1.028 (0.0049) | $2.8 \times 10^{-8}$ |
| 9 | rs11793831 | 23,362,311 | T/G | 0.42 | 1.027 (0.0053) | $4.3 \times 10^{-8}$ |
| 11 | rs1984389 | 31,740,989 | C/A | 0.54 | 0.973 (0.0046) | $2.4 \times 10^{-8}$ |
| 11 | rs10791143 | 131,278,676 | G/A | 0.62 | 1.034 (0.0046) | $1.5 \times 10^{-11}$ |
| 16 | rs4616299 | 7,657,432 | G/A | 0.40 | 0.972 (0.005) | $1.2 \times 10^{-8}$ |
| 17 | rs56058331 | 56,427,128 | A/G | 0.42 | 1.029 (0.0047) | $1.0 \times 10^{-8}$ |
| 18 | rs1261078 | 52,866,791 | G/A | 0.05 | 0.927 (0.0107) | $5.6 \times 10^{-12}$ |
| 19 | rs34232444 | 4,965,404 | C/T | 0.35 | 1.029 (0.0057) | $2.5 \times 10^{-8}$ |
| 19 | rs3746187 | 18,279,816 | G/A | 0.40 | 0.968 (0.0049) | $9.8 \times 10^{-11}$ |
| 19 | rs429358 | 45,411,941 | C/T | 0.15 | 0.942 (0.0067) | $4.6 \times 10^{-19}$ |

**Figure 1.**
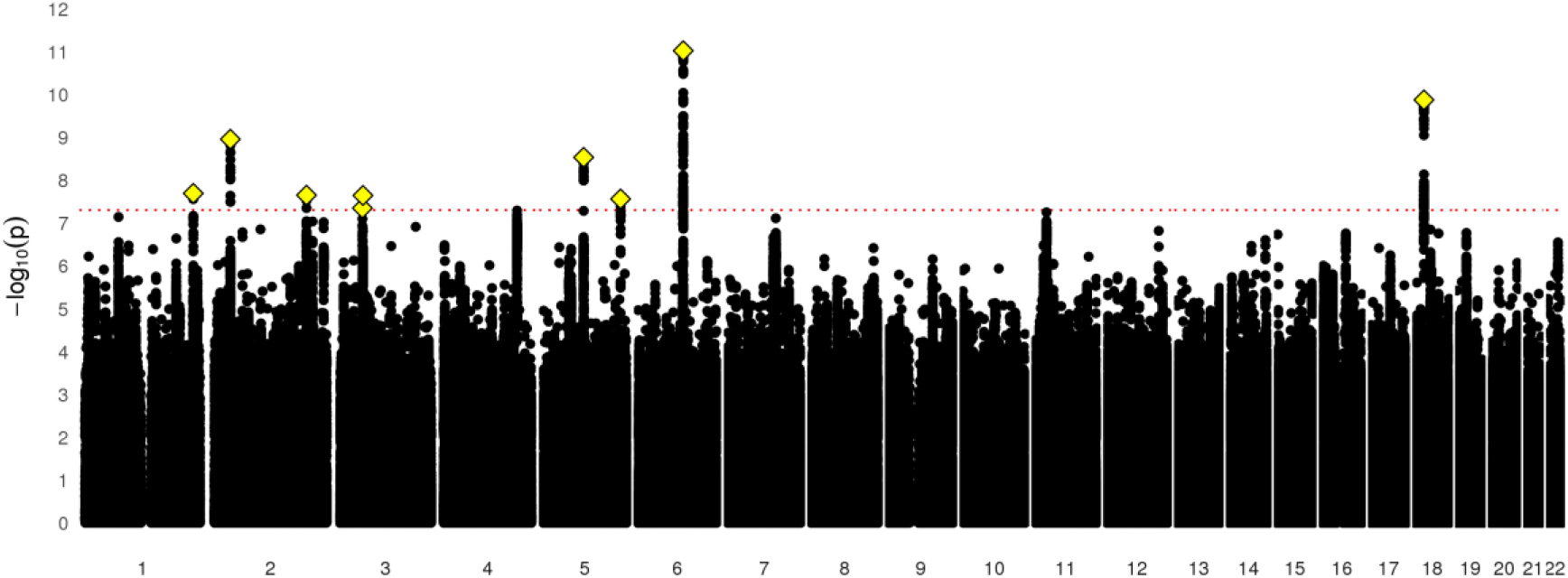
Manhattan plot of email contact in UK Biobank.

**Figure 2.**
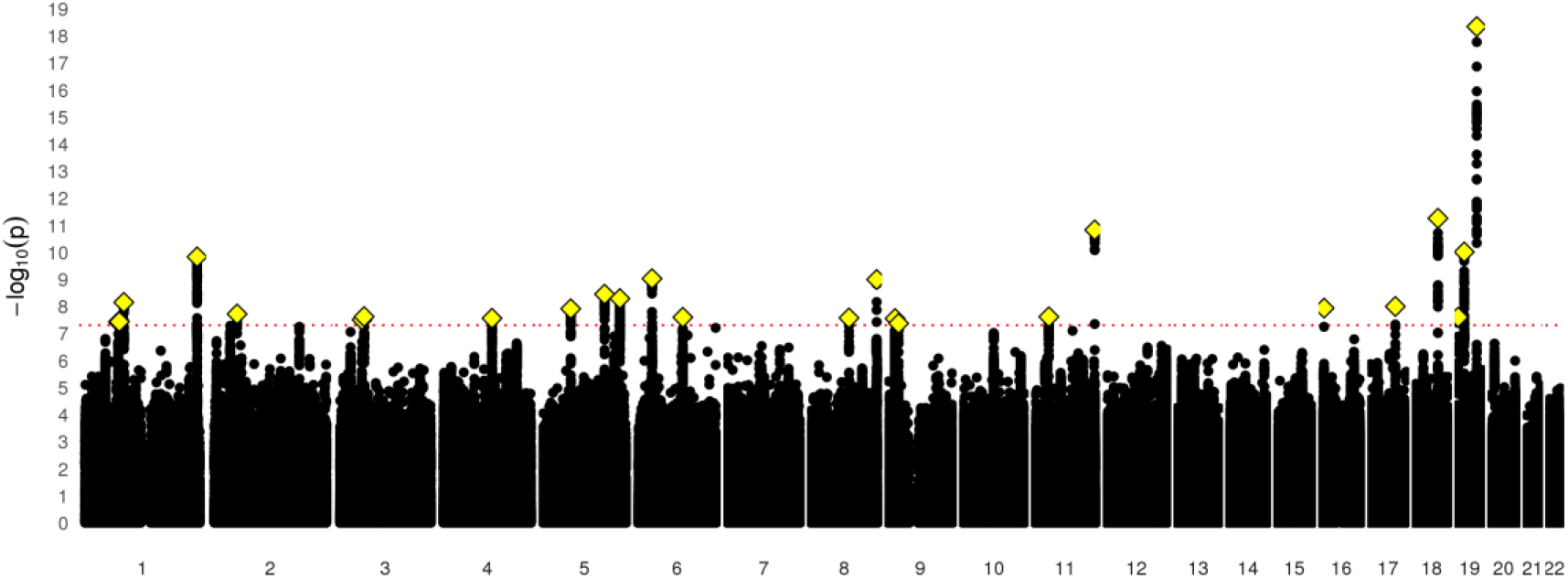
Manhattan plot of data available in MHQ follow-up.

### Loci discovery and annotation of the Email contact and MHQ phenotypes

The nine loci associated with email contact were found to contain an overrepresentation of SNPs found in ncRNA intronic regions (57.5%), as well as SNPs found in intronic regions (28.4%) (Figure 3 and Supplementary Table S1). Evidence was also found that these loci contained regulatory regions of the genome, indicated by 32.0% of the SNPs in the genomic loci having RegulomeDB (RDB) less than 2, indicating that genetic variation in these loci is likely to affect gene expression. Finally, 77.6% of the SNPs within the independent genomic loci had a minimum chromatin state of < 8. This is further evidence that these loci are located in an open chromatin state, providing more evidence that they are located within regulatory regions. Using the GWAS catalogue, lead and tagging SNPs from these 9 independent genomic loci were found to overlap with loci previously associated body mass index and obesity (2 loci), as well as with educational attainment and intelligence (3 loci). (Supplementary Table S2).

**Figure 3.**
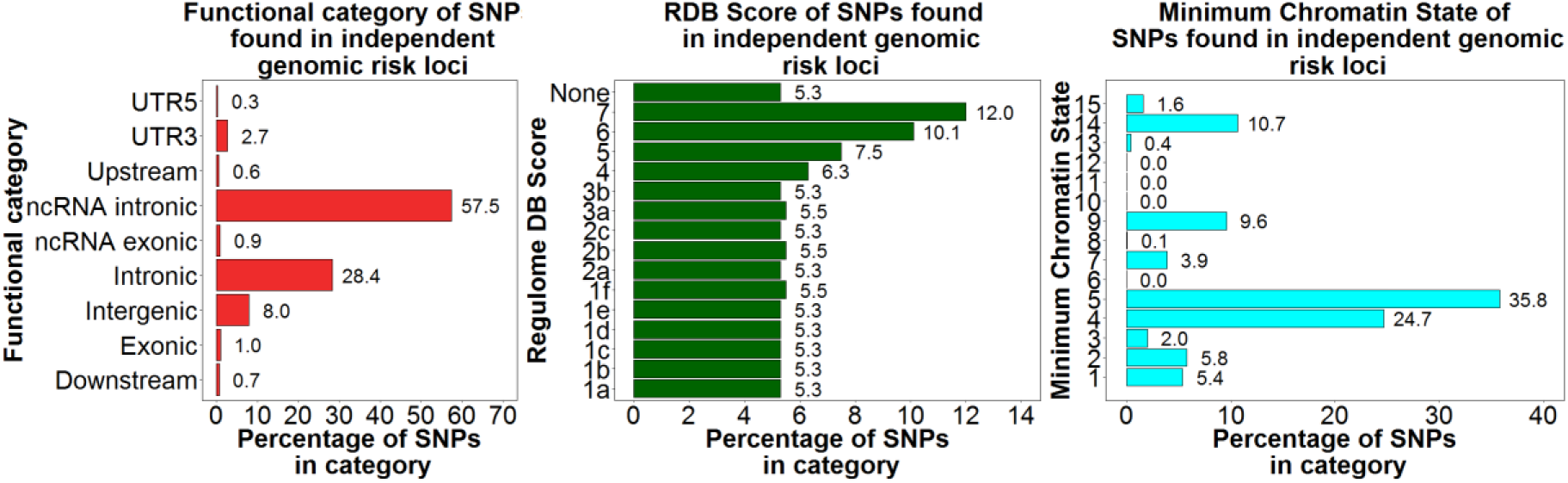
Functional categories, RDB scores, and minimum chromatin states for independent risk loci associated with UKB email contact.

The 25 loci associated with the MHQ participation phenotype notably included rs429358, a missense mutation in *APOE*. The rs429358-C allele is a marker for APOE- ε4 genotype, and the direction of the effect for this SNP indicated that participants with more copies of APOE-ε4 were less likely to participate in the MHQ (OR = 1.029±0.0057SE for each additional ε4 copy). Functional annotation of the SNPs found within these regions showed that these SNPs were primarily located in introns (47.3%), and intergenic regions (17.7%) and 2.9% had no known function (Figure 4 and Supplementary Table S8). Of these SNPs, 30.8% had an RDB score of less than 2 and 83.8% had a minimum chromatin value of less than 8 providing further evidence that these variants are located in regions of the genome that are linked to gene regulation. These 25 loci showed overlap with the loci identified in previous GWAS examining cognitive abilities and education (6 loci), Schizophrenia (5 loci), and Alzheimer’s Disease (1 locus) (Supplementary Table S9).

**Figure 4.**
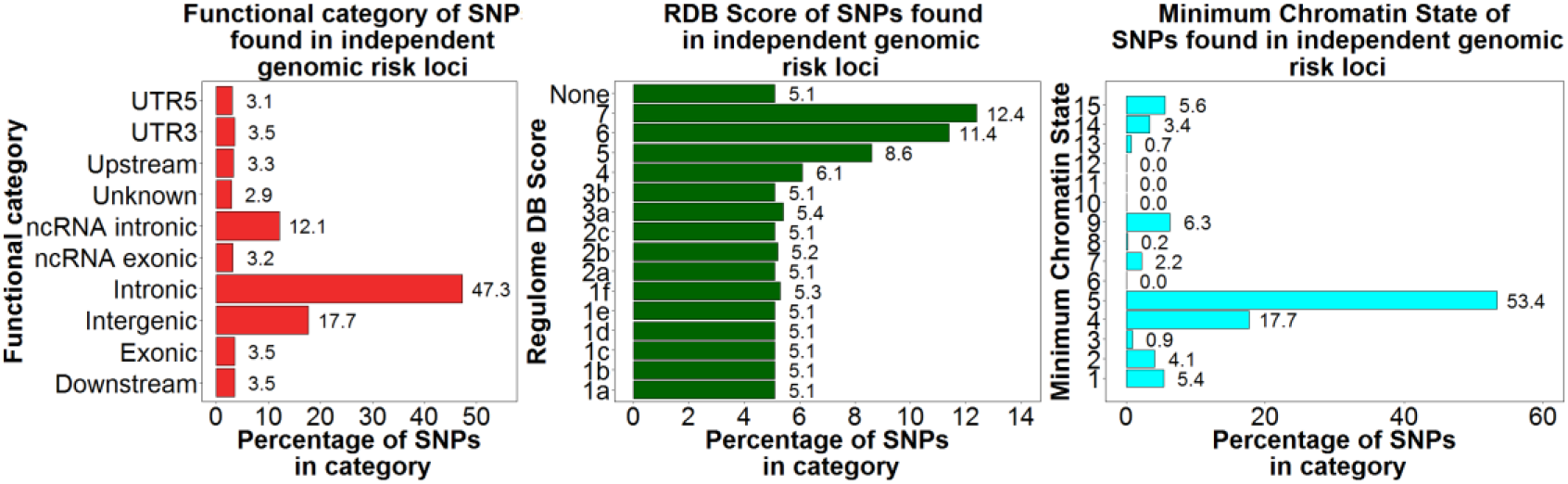
Functional categories, RDB scores, and minimum chromatin states for independent risk loci associated with UKB MHQ participation.

### Gene mapping of the Email access and MHQ phenotype

We used three strategies for mapping the SNPs in the genome wide significant loci to genes. First, positional mapping aligned the SNPs from the independent genomic loci associated with email contact to 20 genes by using location, whereas eQTL mapping matched cis-eQTL SNPs to 40 genes whose level of expression they have been shown to influence. Finally, chromatin interaction mapping annotated SNPs to a total of 41 genes, using three-dimensional DNA-DNA interactions between the SNPs’ genomic regions, and close or distant genes (Supplementary Tables S4 and S5, Supplementary Figure 1a–f). Collectively these mapping strategies identified 70 unique genes, of which 21 were implicated by two mapping strategies and 10 being implicated by all three. A total of five genes, *TNNI3K*, *LRRIQ3*, *NEGR1*, *FPGT*, and *FPGT-TNNI3K*, were implicated using all three methods and showed evidence of a chromatin interaction between two independent genomic risk loci (Supplementary Table S4). Gene-based statistics derived in MAGMA indicated a role for 72 genes (Supplementary Table S5), 4 of which overlapped with genes implicated by all three mapping strategies (Figure 5).

**Figure 5.**
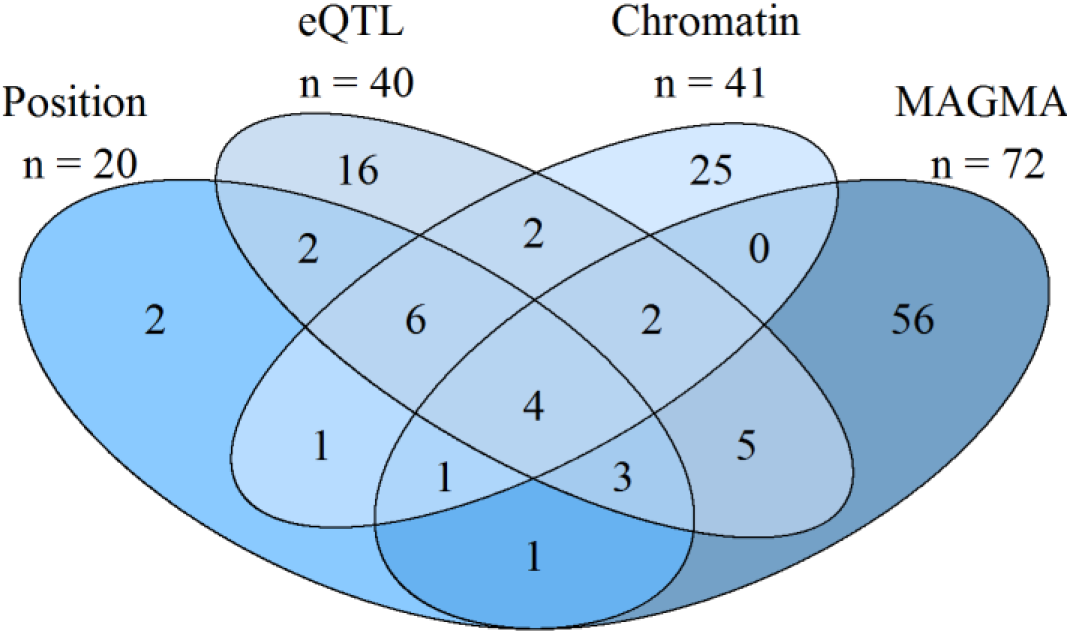
Number of genes implicated by different mapping strategies for UKB email contact.

For the MHQ data phenotype, positional mapping implicated 42 genes, with eQTL mapping indicating a role for 86 genes. Chromatin interaction mapping annotated a total of 124 genes (Supplementary Tables S14 and S15, Supplementary Figure 2a-m). Across these three mapping strategies, 181 unique genes were identified with 46 of these being implicated by two mapping strategies and 25 being implicated by all three. A total of 181 unique genes were implicated by all three mapping strategies. MAGMA was also used to indicate a role for 81 genes (Figure 6 and Supplementary Table S15). Fifteen of these genes overlapped with those identified using the three mapping strategies.

**Figure 6.**
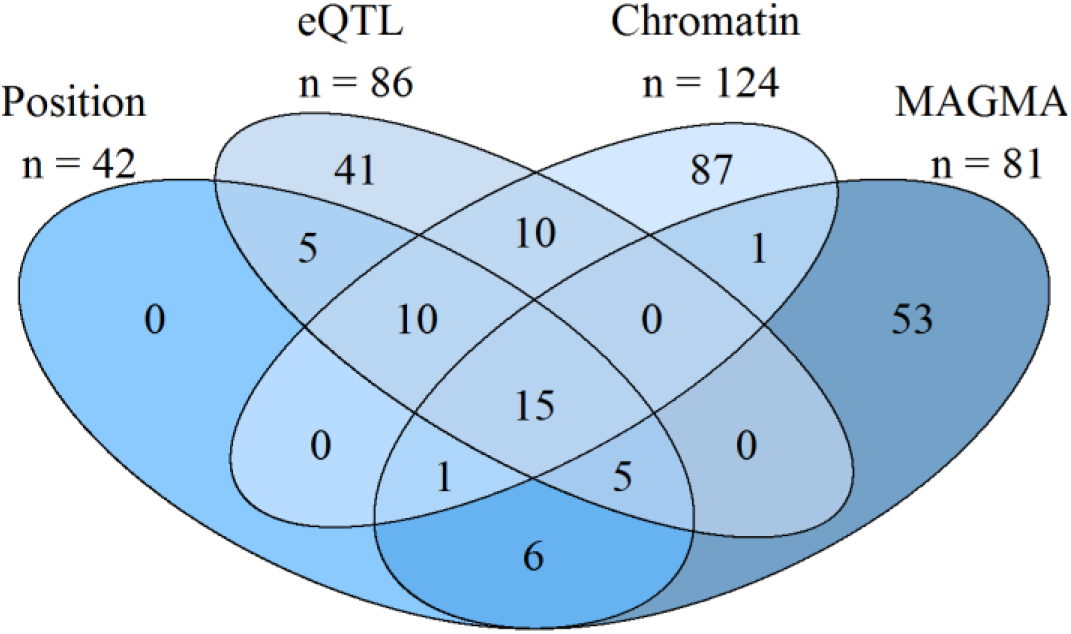
Number of genes implicated by different mapping strategies for UKB MHQ data.

### Gene-set and gene property analysis

The presynaptic membrane gene-set was significantly enriched for the Email contact phenotype (P = 5.19×10^−7^) (Supplementary Table S6). Gene property analysis showed a relationship between expression in the EBV-transformed lymphocyte cells (P = 9.24×10^−4^) and for gene expression in the early mid-prenatal time of life (P = 0.004) (Supplementary Tables S9 and S10).

For the MHQ data phenotype none of the gene sets were enriched (Supplementary Table S16). However, gene property analysis indicated a relationship between gene expression in the brain and the MHQ phenotype (P = 2.64×10^−4^) (Supplementary Table S17) when examining the specific tissue gene groupings this relationship was driven by expression change in the cerebellar hemisphere (P = 8.52×10^−6^) and the Cerebellum (P = 1.27×10^−5^) (Supplementary Table S18). A relationship between gene expression in the early prenatal lifespan range (P = 0.002) and the early mid-prenatal lifespan was also found (P = 5.33×10^−4^) (Supplementary Table S19).

### LD Score regression analysis

We used LD score regression (Bulik-Sullivan et al., 2015) to estimate SNP heritability from the GWAS results. The LD score intercept for email contact and MHQ data in UK Biobank were 1.013 (SE 0.008) and 1.020 (SE 0.008) respectively, while the inflation ratios were 0.037 (SE 0.025) and 0.043 (SE 0.020), respectively. Heritability on the liability scale for email contact was 0.073 (0.004SE) and for MHQ data was 0.099 (0.004SE),. The genetic correlation between email contact and MHQ data was 0.822 (0.020SE).

We used LD Hub (Zheng et al., 2017) to estimate genetic correlations with a large number of other traits. Both email contact and having MHQ data were significantly genetically correlated with a broad spectrum of traits. Results for an illustrative set of traits is plotted in Figure 7 and the results for all traits are listed in Supplementary Table S21. For most anthropometric, behavioral, cognitive, psychiatric, health-related, and life-history traits the direction of the genetic correlations with email contact and MHQ participation was the same. In general, genetic factors associated with providing an email address for recontact to UK Biobank and taking part in the MHQ were also associated with better health, higher intelligence, lower burden of psychiatric disorders, and a slower life-history (e.g., later age at menarche, age at first birth, and menopause). Both email contact and MHQ participation were not significantly genetically correlated with any traits categorized as bone, kidney, uric acid, and metals (transferrin/ferritin). Additionally, email contact was not significantly genetically correlated with glycemic traits while MHQ data availability was not genetically correlated with hormone or metabolite phenotypes.

**Figure 7.**
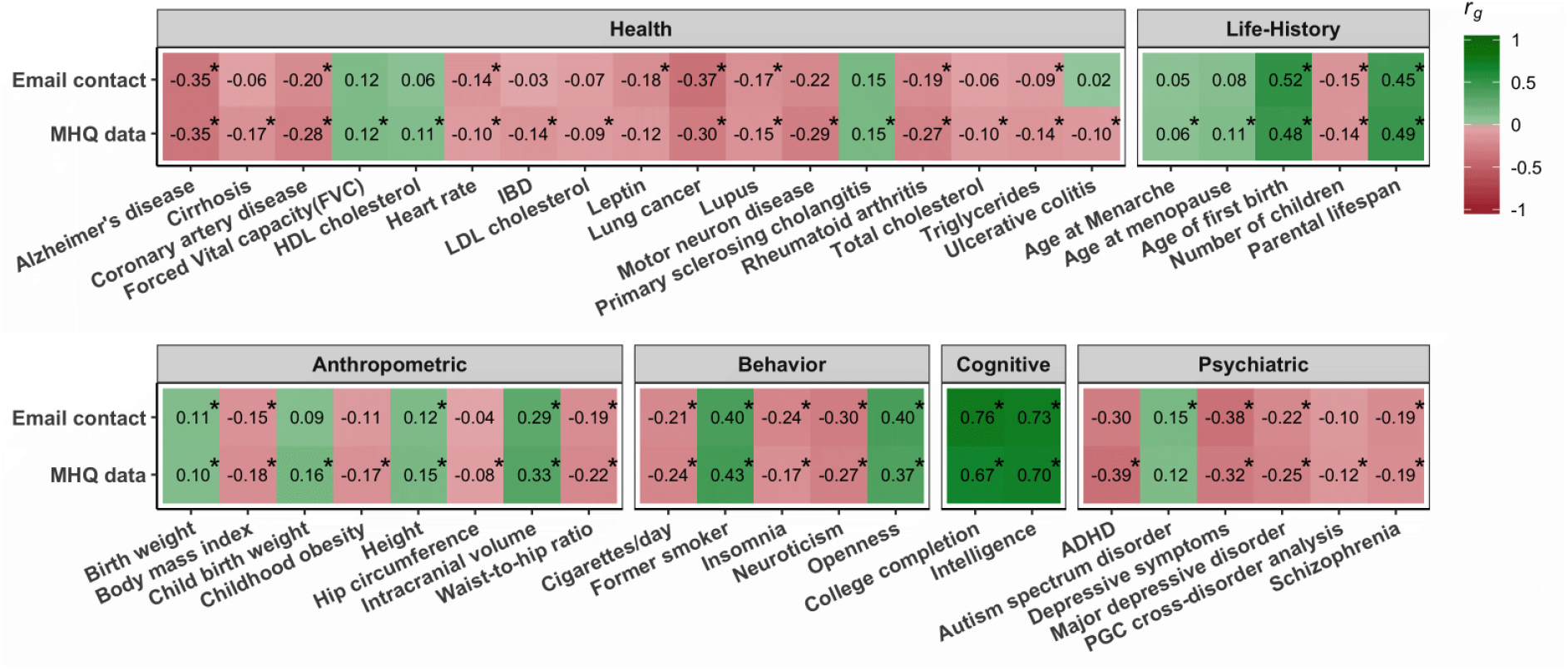
xLD Score genetic correlations (*r_g_*) with email contact and MHQ data. Correlations that are significant at FDR are marked with an asterisk.

### Replication in Generation Scotland

We examined whether any of the associations results for the email and MHQ data phenotypes replicated in an independent sample, using whether members of Generation Scotland participated in the STRADL follow-up of mental health. None of the independent SNPs in the UKB GWASs were significant in Generation Scotland after Bonferroni correction (35 tests) (Supplementary Tables S22 and S23). However, the STRADL data phenotype was genetically correlated with both UKB email contact (*r_g_* = 0.618, p = 1.98 × 10^-6^) and UKB MHQ data (*r_g_* = 0.666, p = 6.12 × 10^-6^) and had a SNP heritability on the liability scale of 0.112 (SE 0.0408).

## Discussion

Using data from UK Biobank, we found that individuals who provided an email address for recontact and who participated in follow-up surveys of mental health differed from those who did not with regards to demographic, psychological, health, and lifestyle, and genetic factors. Most of the phenotypic and genetic associations were in the same direction. These results were not the result of population stratification as only 4% of the inflation in GWAS statistics could be attributed to factors other than polygenic heritability. Having greater educational attainment, being a non-smoker or a former smoker, having fewer hospital diagnoses of illness or injury, and having a family history of dementia or a family history of serious depression all predicted greater likelihood of providing email contact information. Furthermore, in those with that information, those variables were also associated with providing responses to the online Mental Health Questionnaire (MHQ). A few effects went in the opposite direction, with men and younger individuals more likely to provide an email address to UK Biobank, whereas women were more likely to provide MHQ data.

Email contact and MHQ data availability had SNP heritabilities of 7.3% and 9.9% respectively. We identified nine independent SNPs associated with email contact and 25 for MHQ data, more than for many GWAS studies of disease traits in the same sample. Loci for both phenotypes were mostly located within regulatory regions. Of particular interest was the association of MHQ data availability with the apolipoprotein E (APOE) ε4 genotype that is a major risk factor for Alzheimer’s disease. (Coon et al., 2007). While none of these variants individually replicated in an independent data set (Generation Scotland), this may be because Generation Scotland includes a wider age range of participants, the STRADL follow-up was sent by post rather than done online, and because Generation Scotland may be underpowered for finding these effects. However, the strong genetic correlation between STRADL participation and the email contact and MHQ data phenotypes suggests that similar genetic factors are driving participation in follow-up studies.

Email contact and MHQ data shared similar genetic correlations with other traits. There were strong genetic correlations between email contact and indicators of cognitive ability (college completion, *r_g_* = 0.76; intelligence, *r_g_* = 0.73). Contact and data availability were also genetically associated with a *lower burden of genetic risk to mental illness*. The negative genetic correlation with schizophrenia matches results from follow-up participation in the ALSPAC cohort using polygenic risk scores (Martin et al., 2016) but suggests that this association is not specific to schizophrenia.

The similarity in the results for phenotypic and genetic factors associated with email contact and MHQ data show that the availability of an individual to be contacted by email and their choice to participate both act as a filter for selection into the subsample of UK Biobank with Mental Health Questionnaire data. Notably, self-reports of a family history of dementia and a family history of severe depression were more common in email providers and MHQ completers, but individual genetic associations with both these disorders showed significant negative correlations. Individuals who reported dementia or severe depression in their family were therefore more likely to be MHQ participants, even though having a personal genetic predisposition to these disorders may also decrease their likelihood of participating. Knowledge of family history may be a strong motivational factor for participating in follow-up surveys of mental health.

Our sample was large enough that we were able to identify specific genetic loci that were related to participation in follow up studies of mental health. We were also able to analyse the genetics of one particular factor (the availability of email contact for receiving invitations) that is heavily involved in the specific process of follow-up participation. However, a limitation of our analysis is that information on email contact was available for participants at baseline only and thus did not distinguish the entire subset of participants who would have received an email invitation. Another limitation is that information from electronic health records only covered hospital admissions and thus would underestimate associations with milder health conditions.

Individuals in large epidemiological cohorts who participate in follow-up surveys differ in their patterns of phenotypic and genetic association with traits of interest from those who do not. Because most factors had a consistent relationship with the two-step selection process (contactability by email and opting to participate in follow-up), it is likely that these same factors may also differentiate people who choose to become part of the cohort in the first place from other people in the larger population. These factors are very likely to bias the selection of individuals selected for inclusion in population-based studies towards those with positive family histories but lower personal genetic risk of mental health conditions such as depression and dementia. Going forward, studies should evaluate (e.g., using simulations (Munafò et al., 2018)) the particular effects that selection and attrition might have on effect estimates and, where available, check results from follow-up assessments against those from baseline data, even in the cases where the follow-up data provides better or more comprehensive measures of phenotypes of interest. Because continued participation in large cohorts studies recapitulates the “healthy volunteer” effect, comparing responders and non-responders in follow-up surveys may be a useful way of selection bias may influence the generalizability of findings.

## Acknowledgments

MJA, DMH, and AMMc are supported by MRC Mental Health Data Pathfinder award *(Reference MC_PC_17209)* and the Wellcome Trust Strategic Award “STratifying Resilience and Depression Longitudinally” (STRADL) *(Reference 104036/Z/14/Z)*. Analysis conducted under UK Biobank application 4844. WDH is supported by a grant from Age UK (Disconnected Mind Project). IJD is supported by the Centre for Cognitive Ageing and Cognitive Epidemiology, which is funded by the Medical Research Council and the Biotechnology and Biological Sciences Research Council *(Reference MR/K026992/1)*. KASD and MH are supported by NIHR Biomedical Research Centre at South London and Maudsley NHS Foundation Trust and King’s College London. We thank the participants of UK Biobank and Generation Scotland. This work has made use of the resources provided by the Edinburgh Compute and Data Facility (ECDF) (http://www.ecdf.ed.ac.uk/).

