## Supplementary Figures for "Factors associated with sharing email information and mental health survey participation in large population cohorts"

**Supplementary Figures 1 and 2.**

Circos plots by chromosome illustrating genome-wide significant loci associated with the Email contact and the MHQ data phenotype are shown. For each phenotype the most outer layer shows the Manhattan plot and only SNPs where $P < 0.05$ are shown. Each of the SNPs in the genomic risk loci are colour coded indicating the maximum $r^2$ with one of the independent significant SNPs in the locus with red indicating the highest $r^2$ and blue the lowest $r^2$(red $r^2>0.8$, orange $r^2>0.6$, green $r^2>0.4$, and blue $r^2>0.2$). SNPs shown in grey are not in LD with any of the genome wide significant SNPs. The rsID of the most significant lead SNP in each loci is shown. The second layer is the chromosomal ring with the independent genomic risk loci highlighted in blue. Next, the genes mapped by chromatin interactions or eQTLs are displayed. Genes mapped using chromatin interactions the gene is displayed in orange, with genes mapped by eQTL shown in green. Genes that are displayed in red are those mapped using both chromatin interactions and eQTLs. Chromatin interaction links (coloured orange for chromatin interactions and green for eQTLs are displayed.

Figure 1a. Circos plot for email contact chromosome 1

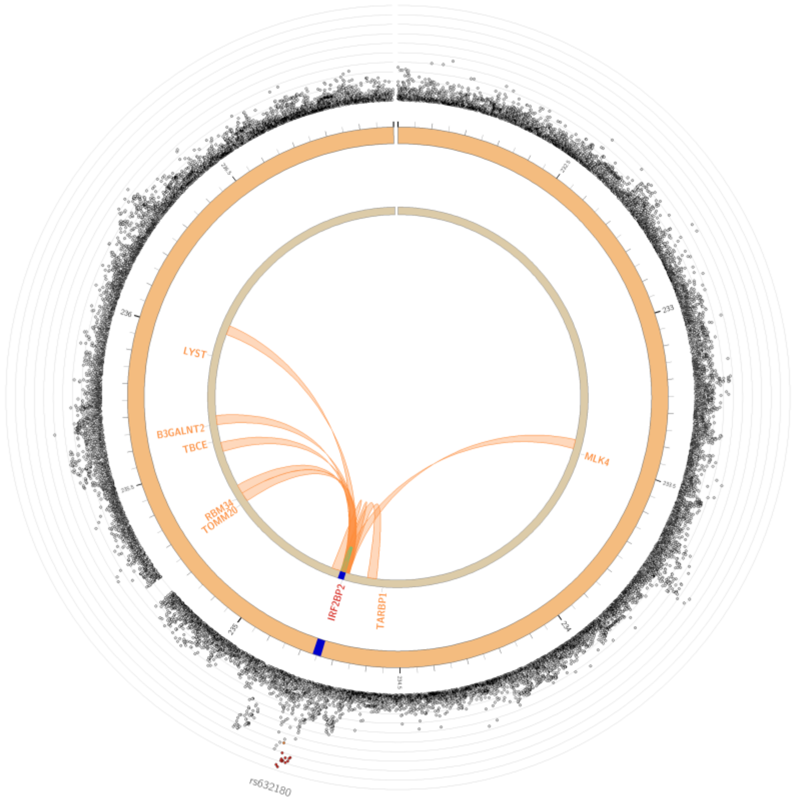


Figure 1b. Circos plot for email contact chromosome 2

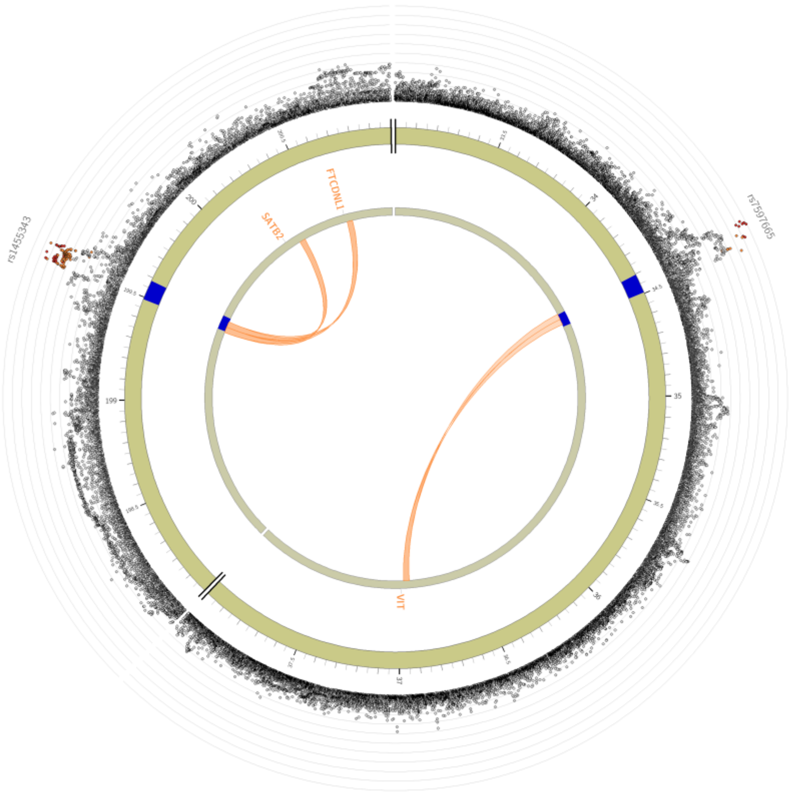


Figure 1c. Circos plot for email contact chromosome 3

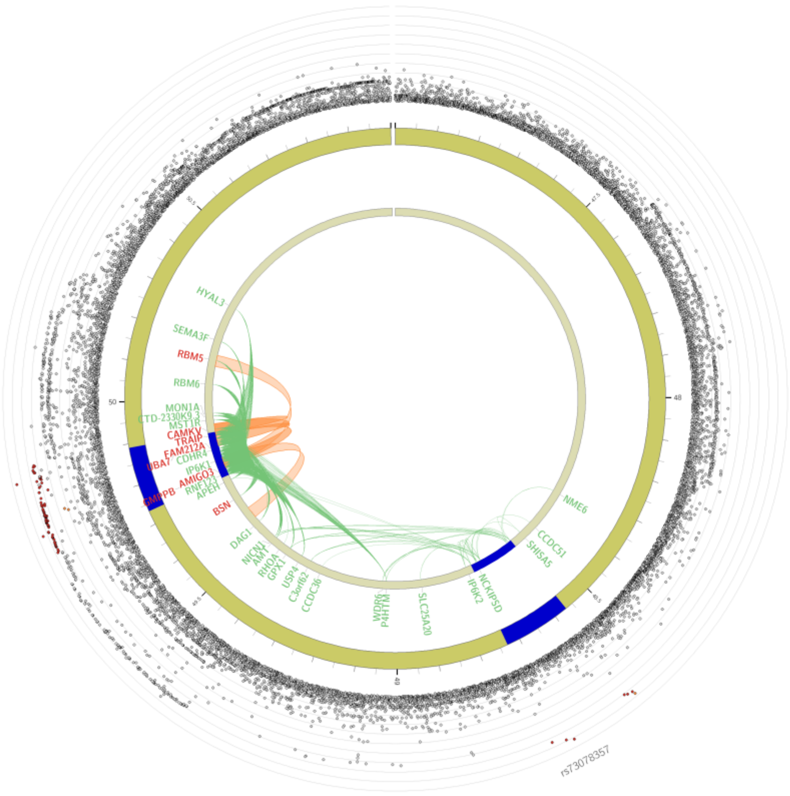


Figure 1d. Circos plot for email contact chromosome 5

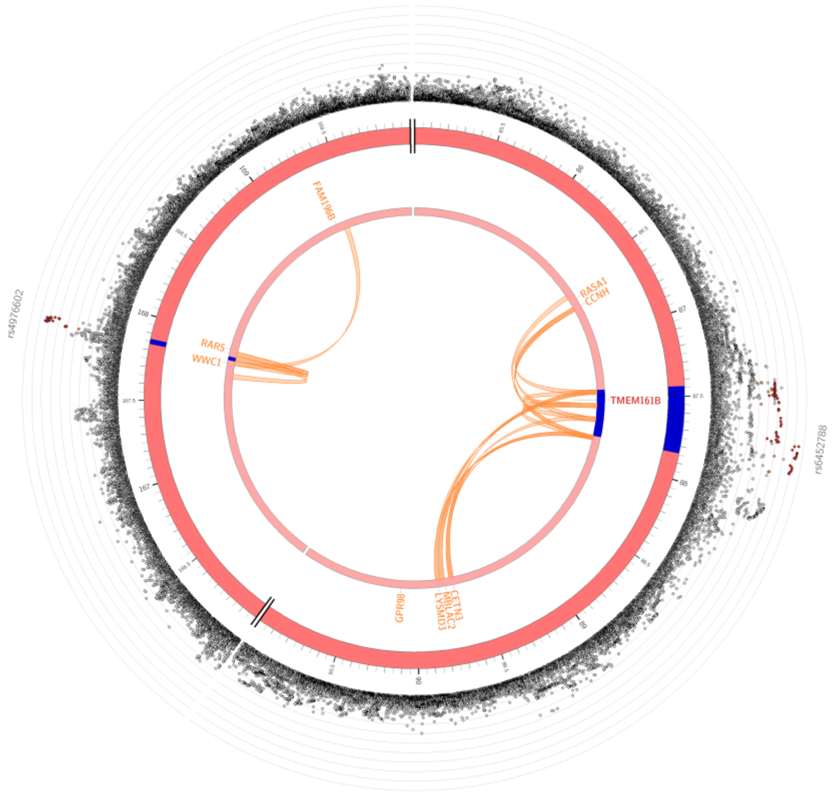


Figure 1e. Circos plot for email contact chromosome 6

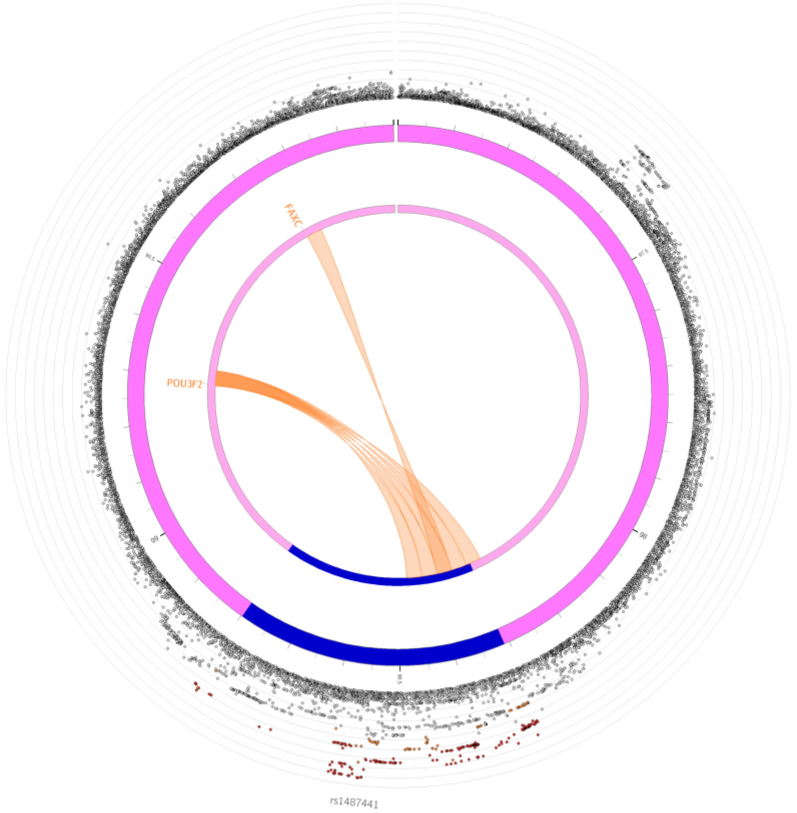


Figure 1f. Circos plot for email contact chromosome 18

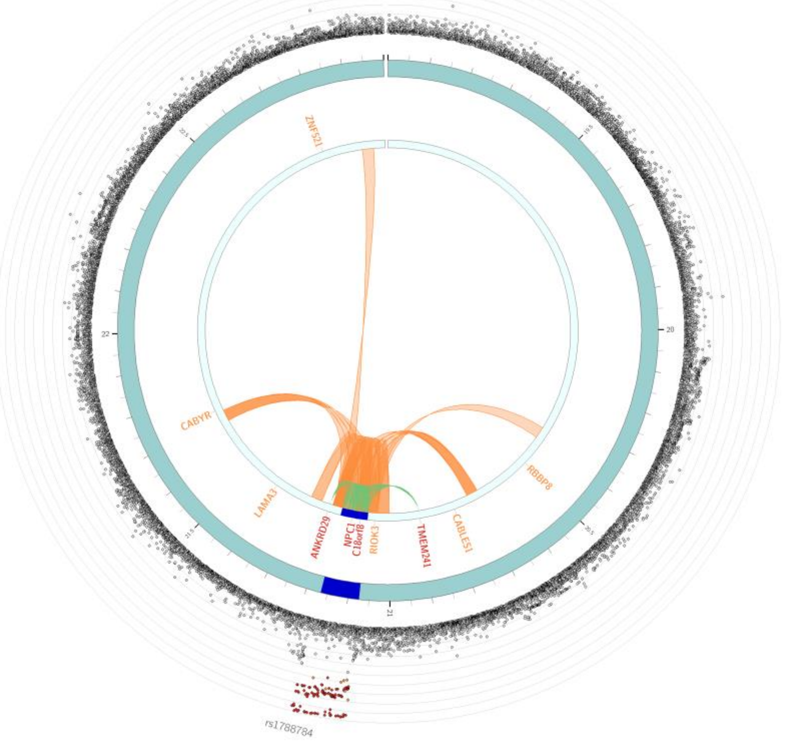


Figure 2a. Circos plot for MHQ data chromosome 1

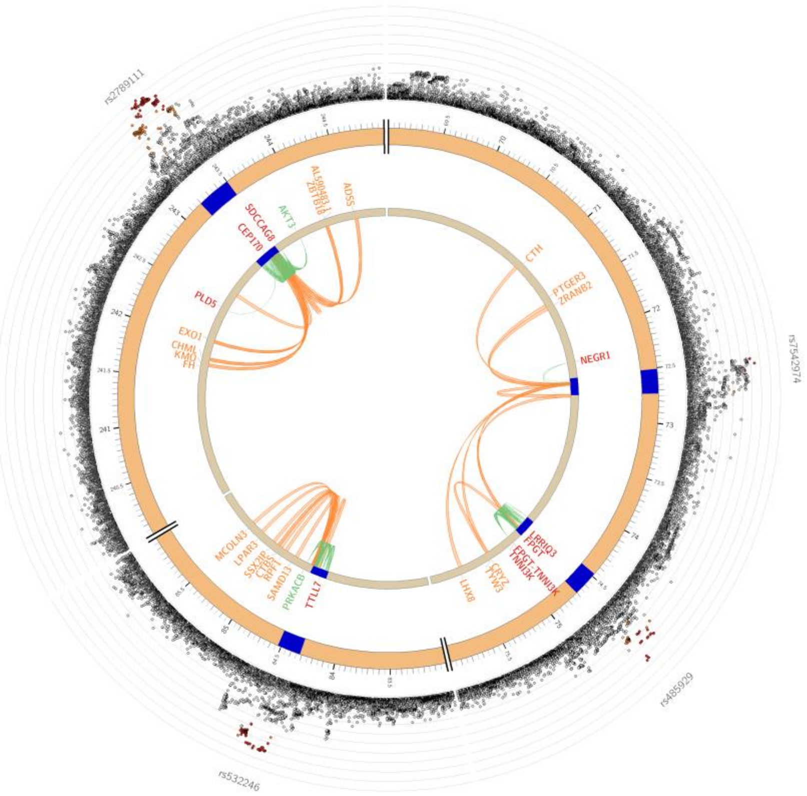


Figure 2b. Circos plot for MHQ data chromosome 2

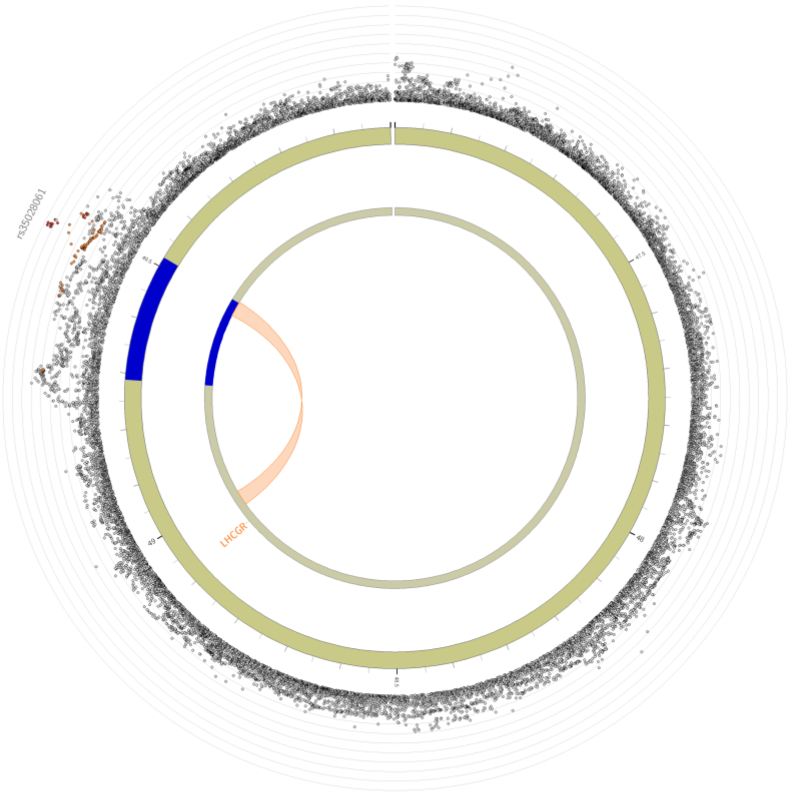


Figure 2c. Circos plot for MHQ data chromosome 3

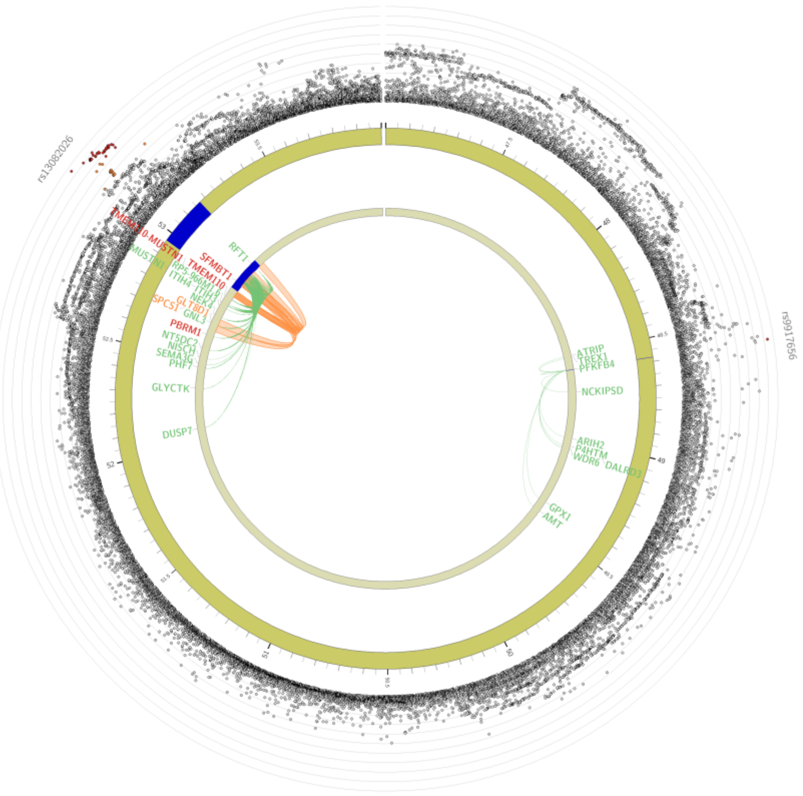


Figure 2d. Circos plot for MHQ data chromosome 4

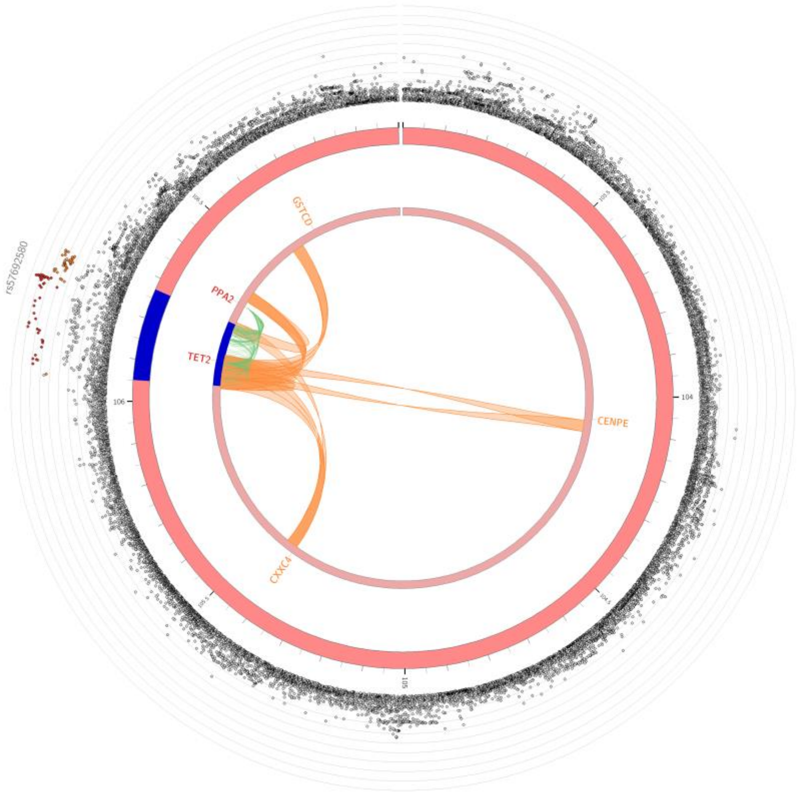


Figure 2e. Circos plot for MHQ data chromosome 5

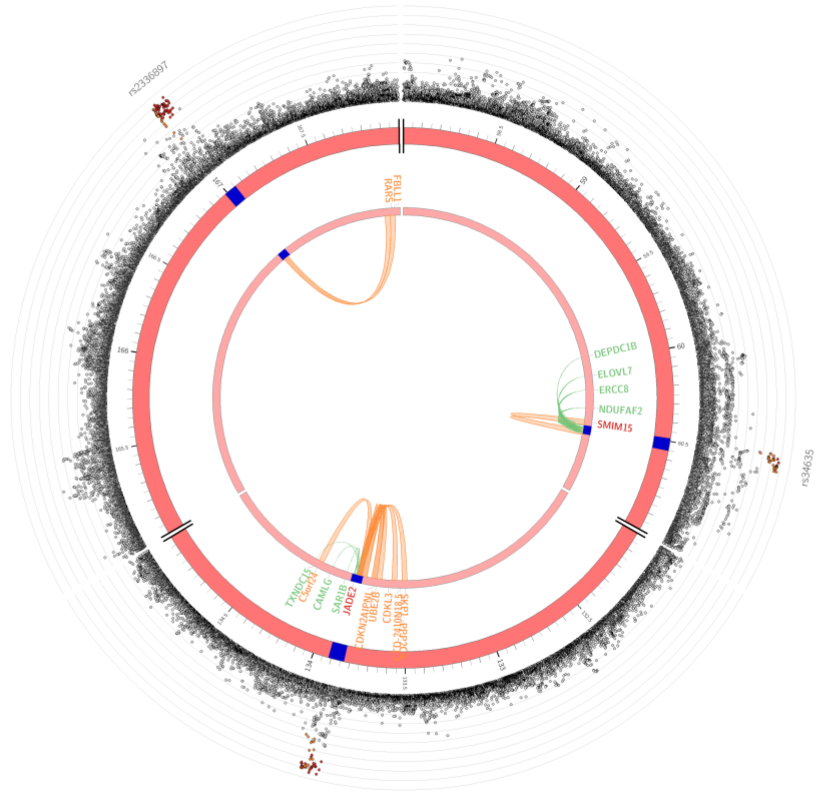


Figure 2f. Circos plot for MHQ data chromosome 6

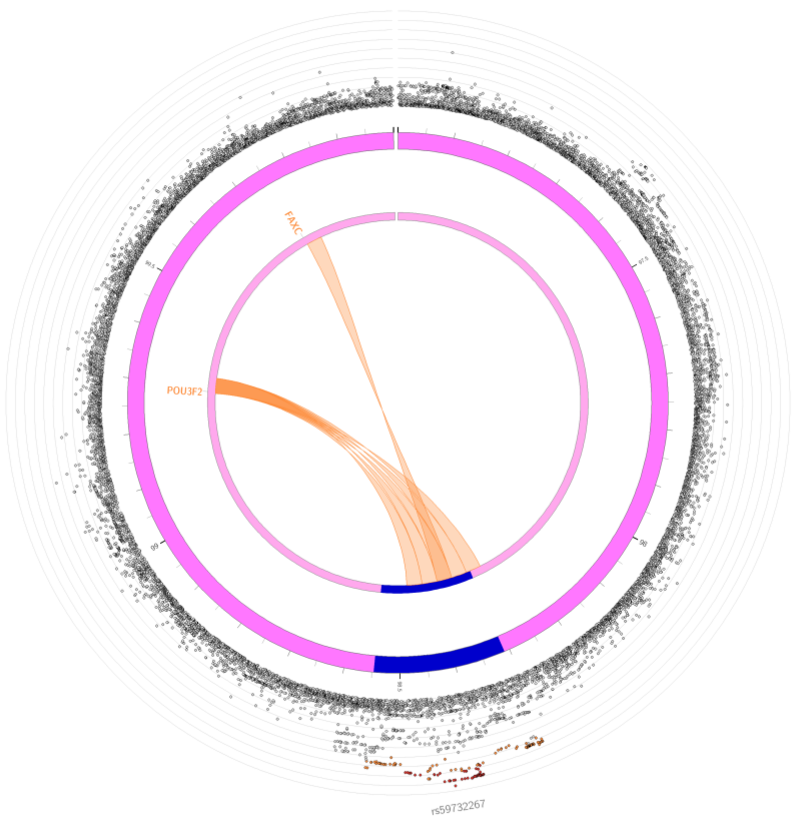


Figure 2g. Circos plot for MHQ data chromosome 8

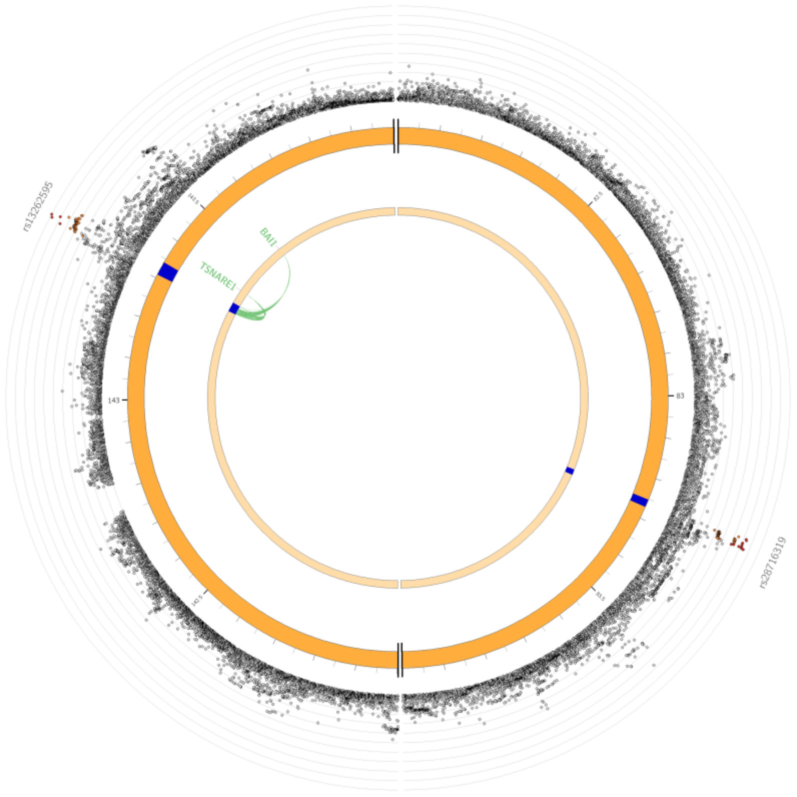


Figure 2h. Circos plot for MHQ data chromosome 9

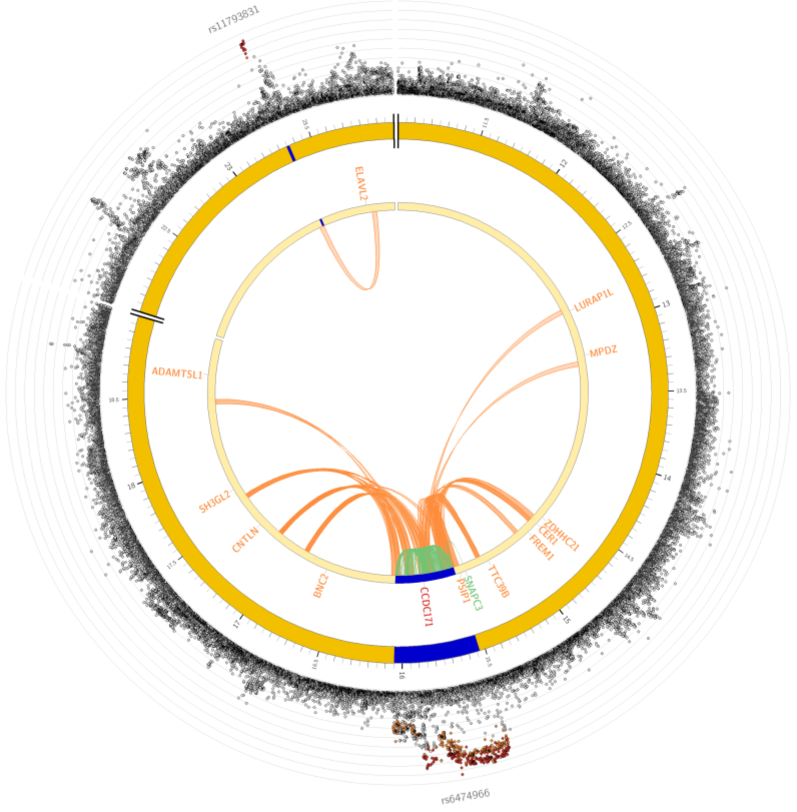

Figure 2i. Circos plot for MHQ data chromosome 11

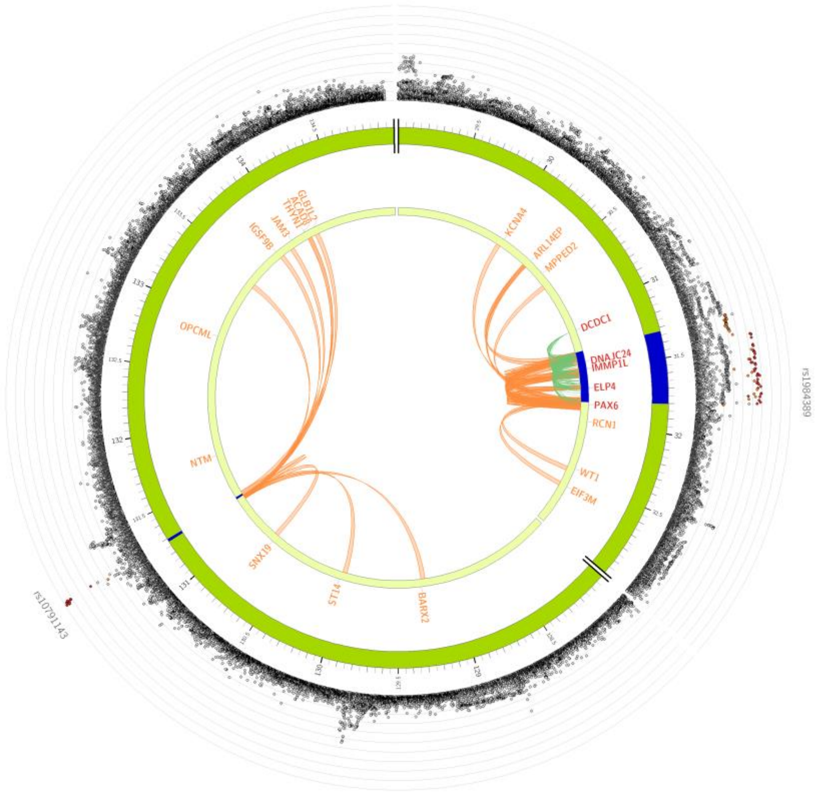

Figure 2j. Circos plot for MHQ data chromosome 16

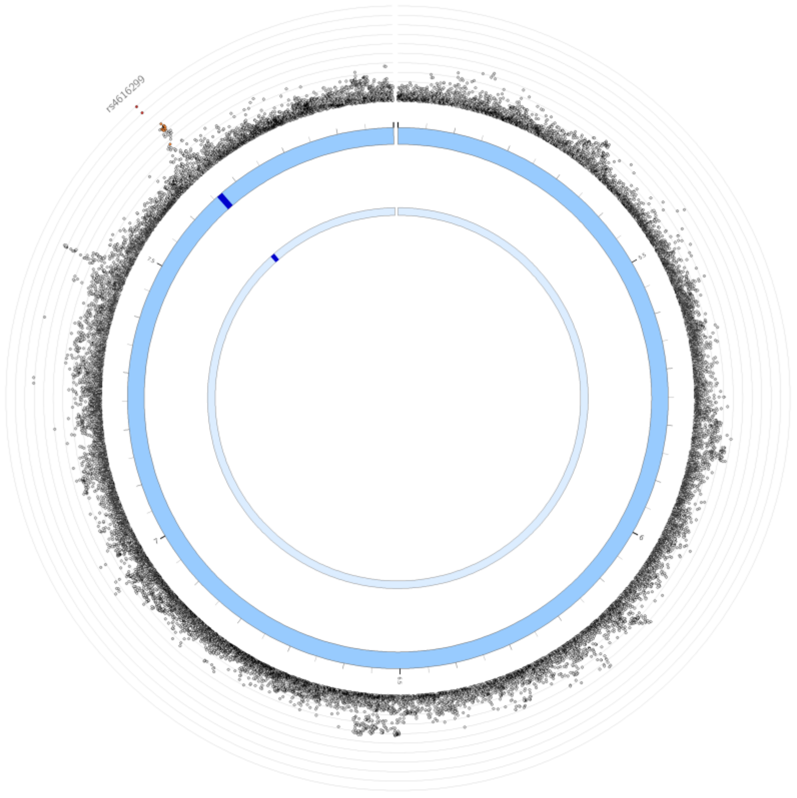


Figure 2k. Circos plot for MHQ data chromosome 17

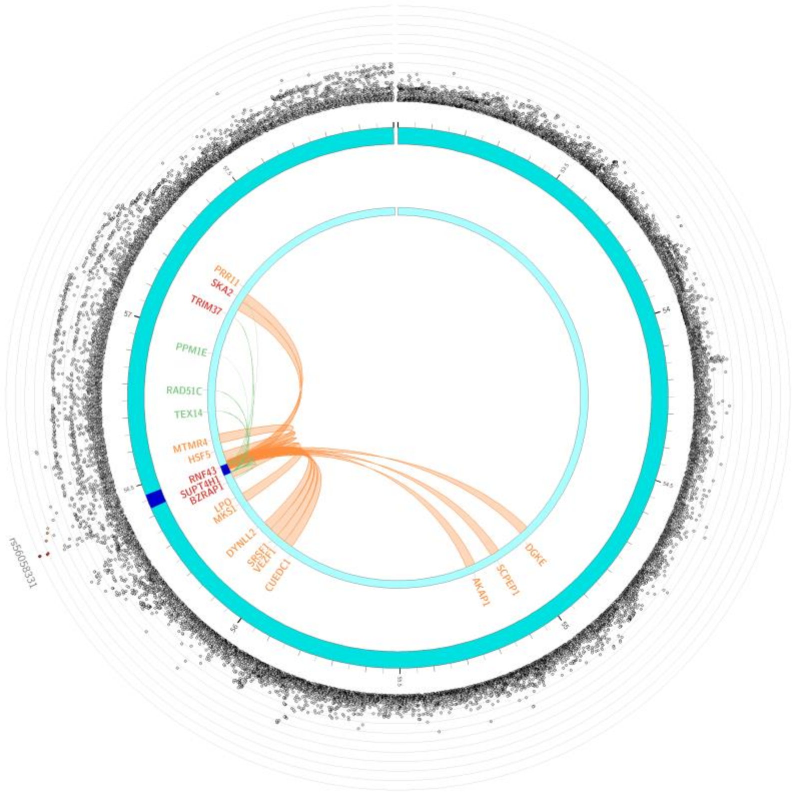


Figure 2l. Circos plot for MHQ data chromosome 18

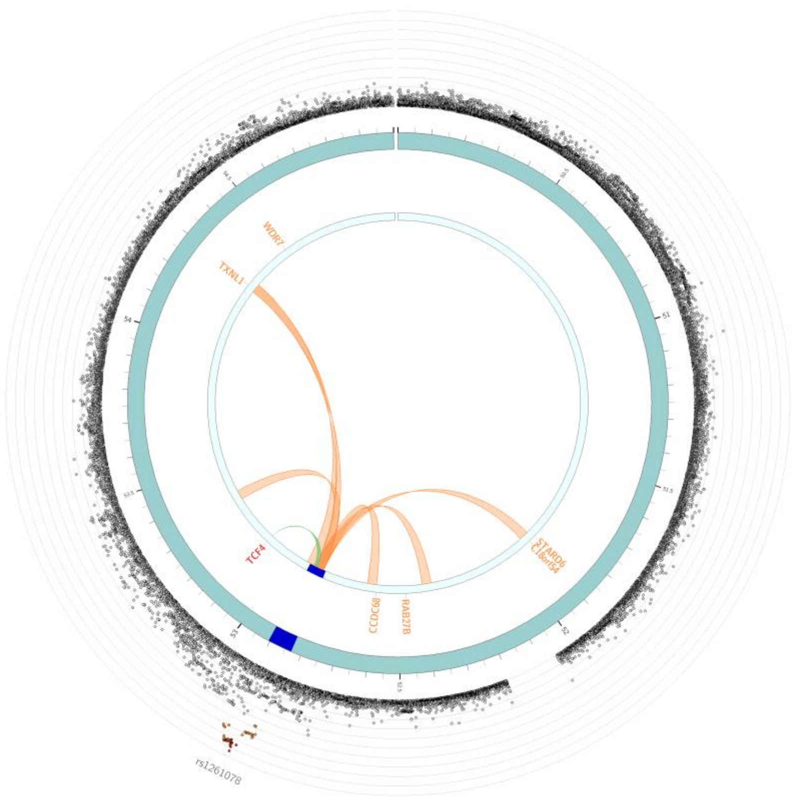


Figure 2m. Circos plot for MHQ data chromosome 19

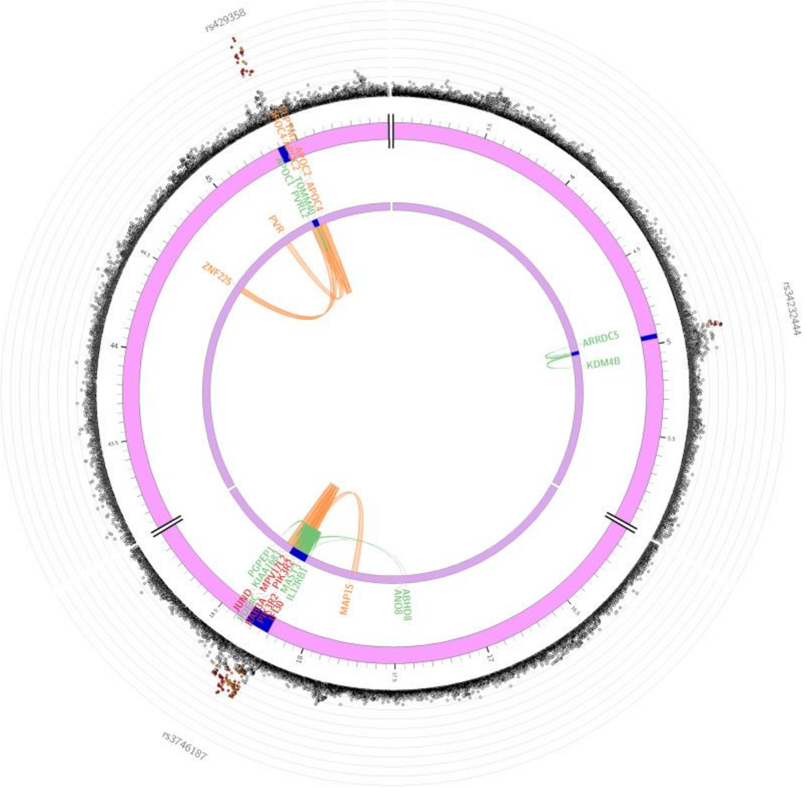

rs429358
rs34232444
rs3746187
ARRDC5
KDM4B
PVR
ZNF225
MAP1S
ABHD8
ANO8
